# Exploring tRNA gene cluster in archaea

**DOI:** 10.1101/370668

## Abstract

Shared traits between prokaryotes and eukaryotes are helpful in the understanding of the tree of life evolution. In bacteria and eukaryotes, it has been shown a particular organization of tRNA genes as clusters, but this trait has not been explored in archaea domain. Here, based on analyses of complete and draft archaeal genomes, we demonstrated the prevalence of tRNA gene clusters in archaea. tRNA gene cluster was identified at least in three *Archaea* class, *Halobacteria*, *Methanobacteria* and *Methanomicrobia* from Euryarchaeota supergroup. Genomic analyses also revealed evidence of tRNA gene cluster associated with mobile genetic elements and horizontal gene transfer inter/intra-domain. The presence of tRNA gene clusters in the three domain of life suggests a role of this type of tRNA gene organization in the biology of the living organisms.

## Introduction

The three domains of life, Archaea, Bacteria and Eukarya are characterized by unique features, even though they share a set of basic characteristics. Archaea and Bacteria, prokaryotes, share ribosomal genes and genes arranged in operons, while archaea and eukaryotes have in common the gene-expression machinery (Reynaud and Devos, 2011). Transfer RNAs (tRNAs) have a major role in the translation machinery and, therefore, belong to the core system of the three domain of life. This molecule is associated with a huge diversity of gene organization, including canonical and disrupted genes. The latter gene species was revealed to be prevalent among archaea genomes (Fujishima and Kanai, 2014).The tRNA genes are found dispersed in the genomes but occasionally, as in mitogenomes, they are organized in clusters containing two to five tRNA genes (Jung et al., 2010; Friedrich et al., 2012; Li et al., 2014). Besides mitogenomes, tRNA gene clusters have been shown to be abundant in Eukarya (Tawari et al., 2008; Bermudez-Santana et al., 2010). In prokaryotes, tRNA gene clusters were characterized in some bacteria, being prevalent in *Firmicutes* phylum and *Mycobacterium* genus (Tran et al., 2016; Morgado and Vicente, 2018), while in archaea, the presence of one tRNA gene cluster was only identified in the *Methanobrevibacter* M1 strain genome (Tran et al., 2016). Considering the common features among the three domains of life we hypothesized the prevalence of tRNA gene cluster also in archaea. In order to test our hypothesis, we analyzed 2481 complete and draft archaea genomes, identifying this structure in Euryarchaeota supergroup, mainly in haloarchaea.

## Material and Methods

### Genomes analyzed

A total of 2481 complete and draft archaeal genomes (available in May 2018) were retrieved from National Center for Biotechnology Information (NCBI) ftp site (ftp://ftp.ncbi.nlm.nih.gov/genomes/genbank/archaea/).

### tRNA gene prediction and tRNA gene cluster identification

The tRNA gene prediction of the data set was performed by tRNAscan-SE 2.0 (Lowe &Chan, 2016). Here we surveyed tRNA gene clusters with a minimum of 10 tRNA genes and used the approach described in a previous study (Morgado and Vicente, 2018) to identify them. tRNA genes were considered clustered if presented a tRNA gene density *≥* 2 tRNA/kb (Bermudez-Santana et al., 2010).

### Genetic analysis

The genetic relationships of the genomes harboring tRNA gene clusters (Euryarchaeota supergroup) were assayed performing a core genome Multilocus Sequence Analysis (cgMLSA). The orthologous genes were retrieved using GET_HOMOLOGUES v3.0.5 (Contreras-Moreira and Vinuesa, 2013) considering a coverage of ≥ 70% and identity ≥ 40%. The 66 orthologous genes were concatenated (yielding ~75 kb length) and submitted to Maximum-likelihood analysis using PhyML v3.1 (Guindon et al., 2010). Genetic relationships of the tRNA genes from the tRNA gene clusters were also assessed by concatenation of their nucleotide sequence (~3 kb length), and subsequent Maximum-likelihood analysis. The alignments were performed by MAFFT v7.271 (Katoh and Standley, 2013), and the tree figures generated by iTOL (Letunic and Bork, 2016).

### Sequence annotation, plasmid and viral identification

The sequence of tRNA gene cluster regions and their flanking regions (~10 kb) were annotated using Prokka v1.12 (Seemann, 2014). A bipartite network of gene content was build using AcCNET v1.2 (Lanza et al, 2017) and visualized in Cytoscape v3.6.0 (Shannon et al, 2003). The topology of the contigs harboring tRNA gene clusters was assayed as described by Jørgensen et al. (2014). These contigs were also submitted to blastn and virSorter (Roux et al., 2015) analysis for plasmid and viral/proviral identification, respectively.

## Results

### Detection and distribution of tRNA gene clusters among archaeagenomes

In order to identify the presence of tRNA gene clusters in archaea genomes we used the methodology described in a previous study (Morgado and Vicente, 2018). The analysis identified tRNA gene clusters in 29/2481 archaea genomes from GenBank (Table 1). The number of tRNA genes in the arrays ranged from 10 to 29, including five clusters with 20 or more tRNA genes. The tRNA clusters encompassed regions from ~1.1 to ~7 kb, representing a density from ~2.14 to ~9.17 tRNA gene/kb (Table 1). The number of tRNA genes present in these clusters corresponds to ~ 17 to 46% of the total number of tRNA genes in these archaea genomes. Two genomes, *Methanobrevibacter* sp. RUG344 and *Methanobrevibacter* sp. RUG833 were the only ones harboring two tRNA gene clusters (Table 1.).

**Table 1.**
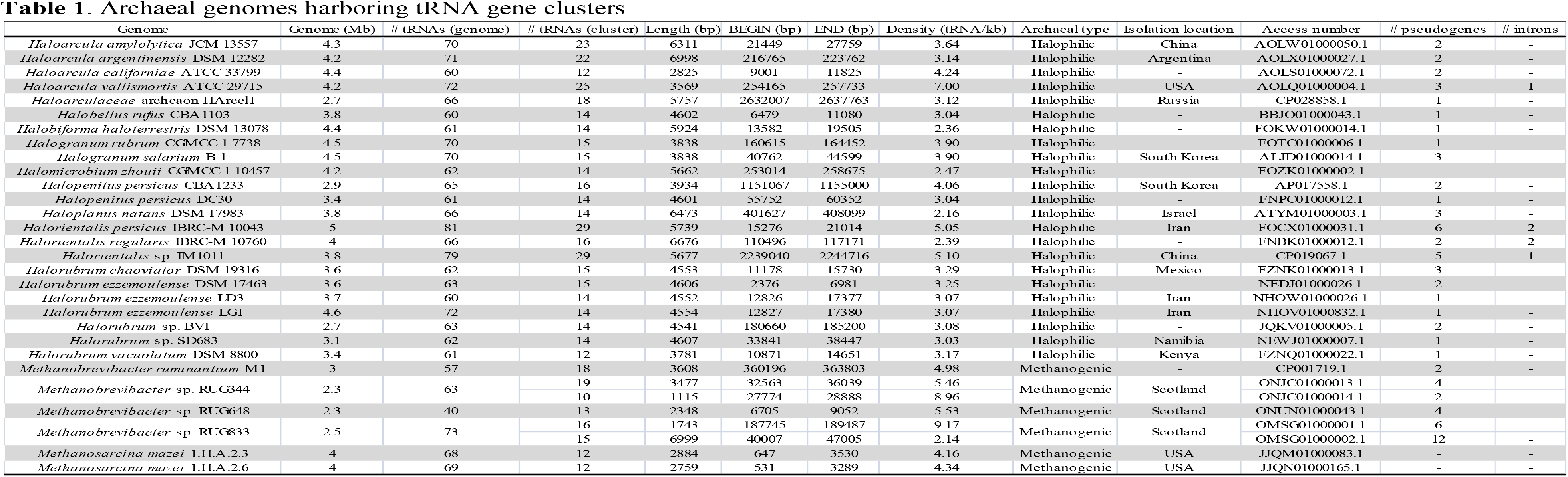
Archaeal genomes harboring tRNA gene clusters

The 29 genomes harboring the tRNA gene clusters correspond to 18 species and seven spp. belonging to 11 archaea genera, all from Euryarchaeota supergroup, including: *Haloarcula*, *Halobellus*, *Halobiforma*, *Halogranum*, *Halomicrobium*, *Halopenitus*, *Haloplanus*, *Halorientalis*, *Halorubrum*, *Methanobrevibacter* and *Methanosarcina*. The latter two genera are from methanogenic archaea, while the others, halophilic. The phylogenetic analysis revealed that all genomes from *Halorubrum* genus but *Halorubrum vacuolatum* DSM 8800 and *Halorubrum* sp. BV1, belong to a single lineage as well as the two genomes from *Methanosarcina mazei* (Fig. 1). Strains harboring tRNA gene clusters with a same tRNA amino acid isotype pattern were isolated from different geographic locations, even considering species from the same lineage (Table 1). Interestingly, 23/29 and 6/29 tRNA gene clusters were identified among 239 halophilic and 618 methanogenic archaea genomes, respectively, revealing a bias to the prevalence of tRNA gene clusters in halophilic archaea. On the other hand, tRNA gene clusters were not identified in any other archaea supergroup.

**Figure 1.**
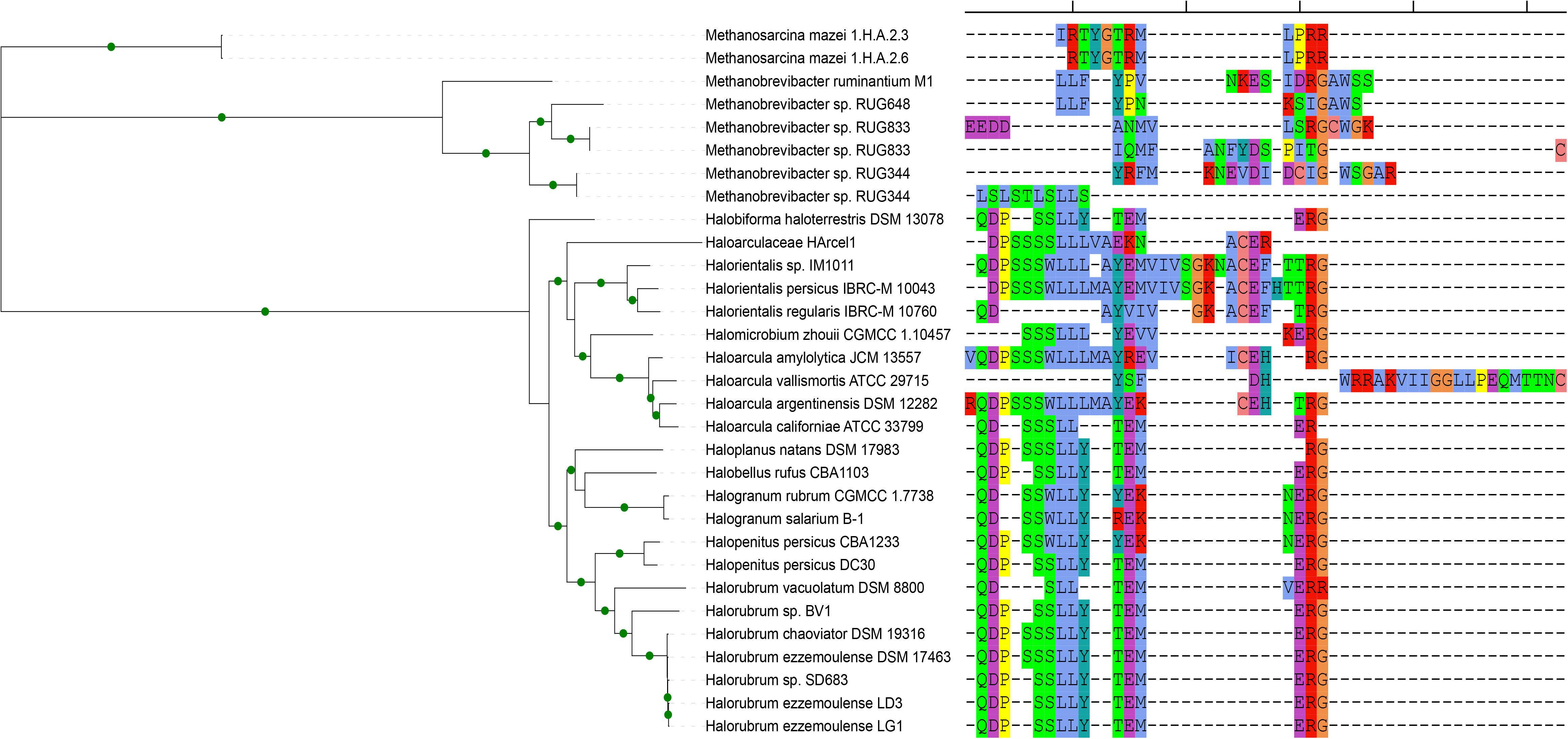
Maximum likelihood tree based on 66 concatenated orthologous genes. In the right side, the tRNA isotype organization (using the single-letter amino acid code) is related with each tree branch. The gaps (- symbol) may not represent the actual distance between two adjacent tRNA genes, but the distance from the reference tRNA gene cluster. The green circles in the branches indicate bootstrap values ≥ 70.

### tRNA gene cluster organization and composition

Overall, the halophilic archaea share tRNA gene clusters presenting similar tRNA amino acid isotype composition and synteny. The exception is the *Haloarcula vallismortis* ATCC 29715 which has a distinct tRNA amino acid isotype composition (Fig. 1). In contrast, the methanogenic archaea harboring tRNA gene clusters present a quite distinct tRNA isotype composition and synteny. This scenario is also observed when considering the nucleotide sequences of the tRNA genes from the clusters (Fig. S1). The most represented isotypes in the tRNA gene clusters were tRNA-Leu and tRNA-Ser, with up to four gene copies per cluster, while the less frequent being tRNA-His and tRNA-Phe, absent in 25 and 21 tRNA gene clusters, respectively (Table S1). Only *Haloarcula vallismortis* ATCC 29715 tRNA gene cluster presented all universal 20 isotypes, while *Halorientalis* sp. IM1011 cluster only has tRNA-His absent. In 26/29 archaea genomes, the tRNA-Tyr and tRNA-Glu from tRNA gene clusters represent a significant mean increment (~45%) to the strains. In general, all tRNA isotypes present in the tRNA gene clusters are redundant with the tRNA isotype repertoire from archaea genomes, the exception is tRNA-Gly in *Methanobrevibacter* sp. RUG833, which is only present in its tRNA gene cluster (Table S1). However, we observed for 9/29 genomes an increment of isoacceptors provided by some tRNA gene clusters, which increased from one to eight species, depending on the strain (Table S2). Almost all clusters presented tRNA genes annotated as pseudogenes, being noteworthy, in *Methanobrevibacter* sp. RUG833, which harbors two tRNA gene clusters summing 31 tRNA genes, being 17 annotated as pseudogenes (Table 1). It was also observed among *Halorientalis* genomes that some tRNA genes in the clusters presented introns.

### Gene content of archaea tRNA gene cluster

The amount of protein-coding genes accompanying the tRNA gene clusters is variable, most of them coding hypothetical proteins. A gene coding for a TROVE domain-containing protein (Bateman and Kickhoefer, 2003) is presented in 20/29 clusters (Fig. S2). The majority of the genes shared by the clusters belong to the halophilic archaea (except for *Haloarcula vallismortis* and *Halogranum* genomes), therefore this group represents the basis of the tRNA gene content network in archaea (Fig. S3).

Several genes associated with DNA transfer mechanisms were identified in the genomic context of the tRNA gene clusters, including transposases, integrases, HNH endonucleases, relaxase, toxin-antitoxin systems and secretion systems (Table S3). Besides these, there were also resistance genes to antibiotics/metals (teicoplanin, bleomycin, chloramphenicol, tetracycline, arsenic). Interestingly, *Halogranum salarium* and *Halogranum rubrum* genomes presented a gene coding for a conjugal transfer protein (HerA). In the methanogenic archaea genomes it was identified a gene coding for a putative phage holin family protein (*Methanobrevibacter ruminantium* M1) and in *Methanosarcina mazei* there were protein-coding genes with high similarity to bacteria, bacteriophages and plasmids from *Proteobacteria* phylum (Table S4), including chloramphenicol O-acetyltransferase gene sequences (Fig. S4).

### tRNA gene clusters in archaea mobilome

Since many genes surrounding the tRNA gene clusters are related to lateral gene transfer and the GC content of some tRNA gene cluster regions contrast with the genomic GC content (Table S5), we surveyed the contigs in order to find sequences related to mobile genetic elements. Among the methanogens, a quite discrepancy intra and inter tRNA gene cluster region GC content could be observed, but none evidence of mobile genetic elements presence was obtained.

By the other hand, in the *Haloarcula californiae* ATCC 33799 strain there is a ~51 kb length contig (access number AOLS01000072.1) harboring the tRNA gene cluster and protein-coding genes with amino acid sequence similarity to a Relaxase (identity 94% and coverage 100%) and a conjugative TraD proteins (identity 95% and coverage 100%) present in the pNYT1 plasmid from *Haloarcula taiwanensis* strain Taiwanensis. In addition, it also presents a type II toxin-antitoxin system from hicA/B family. In *Halorubrum sp.* BV1, a contig with ~188 kb (access number JQKV01000005.1) also presented similarity with sequences from many archaea plasmids, showing coverage of 30% and identity of 96% with *Halorubrum trapanicum* plasmid pCBA1232-02. Besides that, *in-silico* analysis determined the circular topology to this contig, characterizing it as a replicon. Considering all these evidences, it seems that some archaea plasmids can be carrying tRNA gene clusters. Interestingly, some contigs harboring tRNA gene clusters also presented features common to archaea plasmids that could represent an ancestral presence of a plasmid.

Besides the plasmids, we performed a screening for provirus. VirSorter analysis detected putative viral sequences encompassing the regions of the tRNA gene clusters in two genomes, *Haloarcula vallismortis* ATCC 29715 and *Halorientalis* sp. IM101 (Table S6).

## Discussion

Shared traits between prokaryotes and eukaryotes are helpful in the understanding of the tree of life evolution. tRNA gene cluster, a particular feature in RNA gene organization, has been demonstrated to be prevalent in bacteria and eukaryotes, but there is a gap concerning its occurrence in archaea. Although the significant presence among bacteria and eukaryotes, the role of the tRNA gene cluster is under debate. In bacteria, it has been implicated with a faster growing (Ardell and Kirsebom, 2005) and in the modulation of the tRNA transcription and translation process (Rudner et al., 1993). In eukaryotes, there is a negative correlation between clusters of tRNA genes and chromosomal stability, since they can act as barriers to DNA replication and the consequent formation of genomic fragile sites (Bermudez-Santana et al., 2010). Besides that, the tRNA-derived fragments (tsRNAs or tRFs), identified in the three domain of life (Zhu et al., 2018), could be generated from the tRNA gene clusters, being another common trait between prokaryotes and eukaryotes.

In this study, based on analyses of complete and draft archaeal genomes we demonstrated the prevalence of tRNA gene clusters in archaea. Considering the four archaea supergroups (Asgard, DPANN, TACK and Euryarchaeota)(Spang et al., 2017), tRNA gene cluster occurs at least in three class (*Halobacteria*, *Methanobacteria* and *Methanomicrobia*) from Euryarchaeota. In fact, there is a bias concerning the number of genomes and drafts from Euryarchaeota, and therefore it is not possible to assume that this trait is exclusive of this supergroup.

Similarly to what was found in bacteria (Talla et al. 2013; Morgado and Vicente, 2018), archaea tRNA gene clusters favor tRNA isotype redundancy, just increasing the number of tRNA gene copies. Another common trait between archaea and bacteria is the presence of protein coding genes within some tRNA gene clusters. In archaea, most of tRNA gene clusters harbor less than 20 tRNA genes, while in bacteria, there is a considerable number of tRNA gene clusters with more than 20 tRNA genes. Besides tRNA genes, there are also tRNA pseudogenes in the tRNA gene clusters from prokaryotes and eukaryotes (Giroux and Cedergren, 1988; Bermudez-Santana et al., 2010; Puerto-Galan and Vioque, 2012), suggesting a compromised canonical function.

Concerning the origin of the tRNA gene clusters, in bacteria it was assumed that they originated from *Firmicutes* phylum through a horizontal gene transfer event (Talla et al. 2013). Indeed, there are evidence showing the association of the tRNA gene clusters with mobile genetic elements. In several bacteria species there are plasmids carrying tRNA gene clusters (Talla et al. 2013; Morgado and Vicente, 2018), and in archaea we have evidences that some contigs harboring tRNA gene clusters are associated with homologous sequences of archaea plasmids (this work). Interestingly, one of these contigs presented a relaxase gene, which is rare in archaeal plasmids (Smillie et al., 2010; Guglielmini et al., 2013), but a common trait of conjugative bacterial plasmids. Moreover, tRNA gene clusters are abundant in mycobacteriophages (Hatfull et al., 2010; Pope et al., 2014; Delesalle et al., 2016), also occurring in some bacteriophages (Bailly-Bechet et al., 2007). So far, tRNA gene cluster had been only identified in one haloarchaeal virus genome, being composed of 36 tRNA genes (Senčilo et al., 2013). Here, using VirSorter, it was raised an evidence of two archaea genomes presenting tRNA gene clusters in regions predicted as provirus sequences.

All archaea genomes harboring tRNA gene clusters were from Euryarchaeota supergroup, which encompasses some mesophilic strains. In fact, it has been reported events of bacterial gene acquisition by mesophilic archaea, revealing the existence of horizontal gene transfer between these two domains of life (Wagner et al., 2017). Particularly, in the tRNA gene clusters from the mesophilic methanogenic *Methanosarcina mazei* genomes, we identified regions with high similarity to bacteria and bacteriophages, suggesting the bacteria as the source of these genes. In fact, *Caudovirales* order infects both archaea and bacteria (Prangishvili et al., 2006; Iranzo et al. 2016), acting as gene vectors between these domains.

Therefore, it is clear that tRNA gene cluster is a common trait in the three domain of life, that independently has been evolving in each domain.

